# Expectancy-related changes in firing of dopamine neurons depend on hippocampus

**DOI:** 10.1101/2023.07.19.549728

**Authors:** Yuji K. Takahashi, Zhewei Zhang, Marlian Montesinos-Cartegena, Thorsten Kahnt, Angela J. Langdon, Geoffrey Schoenbaum

**Affiliations:** Intramural Research Program, National Institute on Drug Abuse, Baltimore, MD; Intramural Research Program, National Institute on Mental Health, Bethesda, MD

**Author notes:** Correspondence or requests for material should be addressed to Y.T. or G.S. shared first/last authorship (Yuji K. Takahashi, Zhewei Zhang, Angela J. Langdon, Geoffrey Schoenbaum).

**Keywords:** dopamine, hippocampus, prediction error, learning, single-unit, rodent

## Abstract

The orbitofrontal cortex (OFC) and hippocampus (HC) are both implicated in forming the cognitive or task maps that support flexible behavior. Previously, we used the dopamine neurons as a sensor or tool to measure the functional effects of OFC lesions (Takahashi et al., 2011). We recorded midbrain dopamine neurons as rats performed an odor-based choice task, in which errors in the prediction of reward were induced by manipulating the number or timing of the expected rewards across blocks of trials. We found that OFC lesions ipsilateral to the recording electrodes caused prediction errors to be degraded consistent with a loss in the resolution of the task states, particularly under conditions where hidden information was critical to sharpening the predictions. Here we have repeated this experiment, along with computational modeling of the results, in rats with ipsilateral HC lesions. The results show HC also shapes the map of our task, however unlike OFC, which provides information local to the trial, the HC appears to be necessary for estimating the upper-level hidden states based on the information that is discontinuous or separated by longer timescales. The results contrast the respective roles of the OFC and HC in cognitive mapping and add to evidence that the dopamine neurons access a rich information set from distributed regions regarding the predictive structure of the environment, potentially enabling this powerful teaching signal to support complex learning and behavior.

## Introduction

The orbitofrontal cortex (OFC) and hippocampus (HC) are both implicated in learning the states, and relationships between them, that define the world around us (Eichenbaum, 2000; O’Keefe and Nadel, 1978; Wilson et al., 2014). These so-called cognitive or task maps are fundamental to the process by which we predict impending events, particularly valuable outcomes (Behrens et al., 2018; Tolman, 1948). Both areas are also well-positioned to provide information, either directly or indirectly, to the midbrain dopamine neurons, which generate teaching signals that reflect discrepancies between actual and expected outcomes (Mirenowicz and Schultz, 1994; Schultz, 2016; Starkweather and Uchida, 2021).

Several years ago, we capitalized on this arrangement by using the dopamine neurons as a sensor or tool to measure the functional effects of OFC lesions (Takahashi *et al*., 2011). We recorded midbrain dopamine neurons as rats performed an odor-based choice task, in which errors in the prediction of reward were induced by manipulating the number or timing of the expected rewards across blocks of trials. We found that OFC lesions ipsilateral to the recording electrodes caused prediction errors to be degraded. In particular, the response to reward omission was largely abolished, while the response to unexpected reward was both diminished and slower to adapt with learning across trials. Activity to the predictive cues was also less closely tied to high and low values. Critically, these results were not consistent with our hypothesis going in that the OFC provided information about the value of expected outcomes, and instead a computational modeling approach showed it was best understood as a loss in the resolution of the task states, particularly under conditions where hidden information was critical to sharpening the predictions. Notably these data led to the hypothesis that the OFC is a critical part of the cognitive mapping circuit (Wilson *et al*., 2014).

Here we have repeated this experiment, along with computational modeling of the results, to test the hypothesis that HC provides similar information to the dopamine neurons regarding the layout of the task space. The results show that this is indeed the case, however unlike the OFC, which provides information local to the trial, the HC appears to be necessary only for estimating the upper-level hidden states based on the information that is discontinuous or separated by longer timescales. The results provide novel data contrasting the respective roles of the OFC and HC in cognitive mapping and how these areas interact to support learning. Additionally, they add to evidence that the dopamine neurons access a rich information set from distributed regions regarding the predictive structure of the environment, potentially enabling this powerful teaching signal to support complex learning and behavior.

## Results

We recorded single-unit activity in the VTA of rats with ipsilateral sham (n = 5) or neurotoxic (n = 9) lesions targeting the HC and resulting in visible loss of neurons in 54% (49-60%) of this region across subjects (Figure S1). Single-unit activity was recorded as rats performed an odor-guided choice task identical to the one we previously used to characterize signaling of reward predictions and reward prediction errors in rats with ipsilateral OFC lesions (Takahashi *et al*., 2011). On each trial, rats sampled one of three different odor cues at a central port and then responded at one of two adjacent fluid wells (Fig. 1a). One odor signaled the availability of sucrose reward only in the left well (forced left), a second odor signaled sucrose reward only in the right well (forced right), and a third odor signaled the reward was available at either well (free choice). To induce errors in reward prediction, we manipulated either the timing or the number of rewards delivered in each well across 4 blocks of trials (Fig. 1b). Positive prediction errors were induced by making a previously delayed reward immediate (Fig. 1a, 2^*sh*^ and 1^*st*^ bolus in 3^*bg*^) or by adding more reward (Fig. 1b, 2^*nd*^ bolus in 3^*bg*^ and 4^*bg*^), whereas negative prediction errors were induced by delaying a previously immediate reward (Fig. 1b, 2^*lo*^) or by decreasing the number of reward (Fig. 1b, 4^*sm*^).

**Figure 1:**
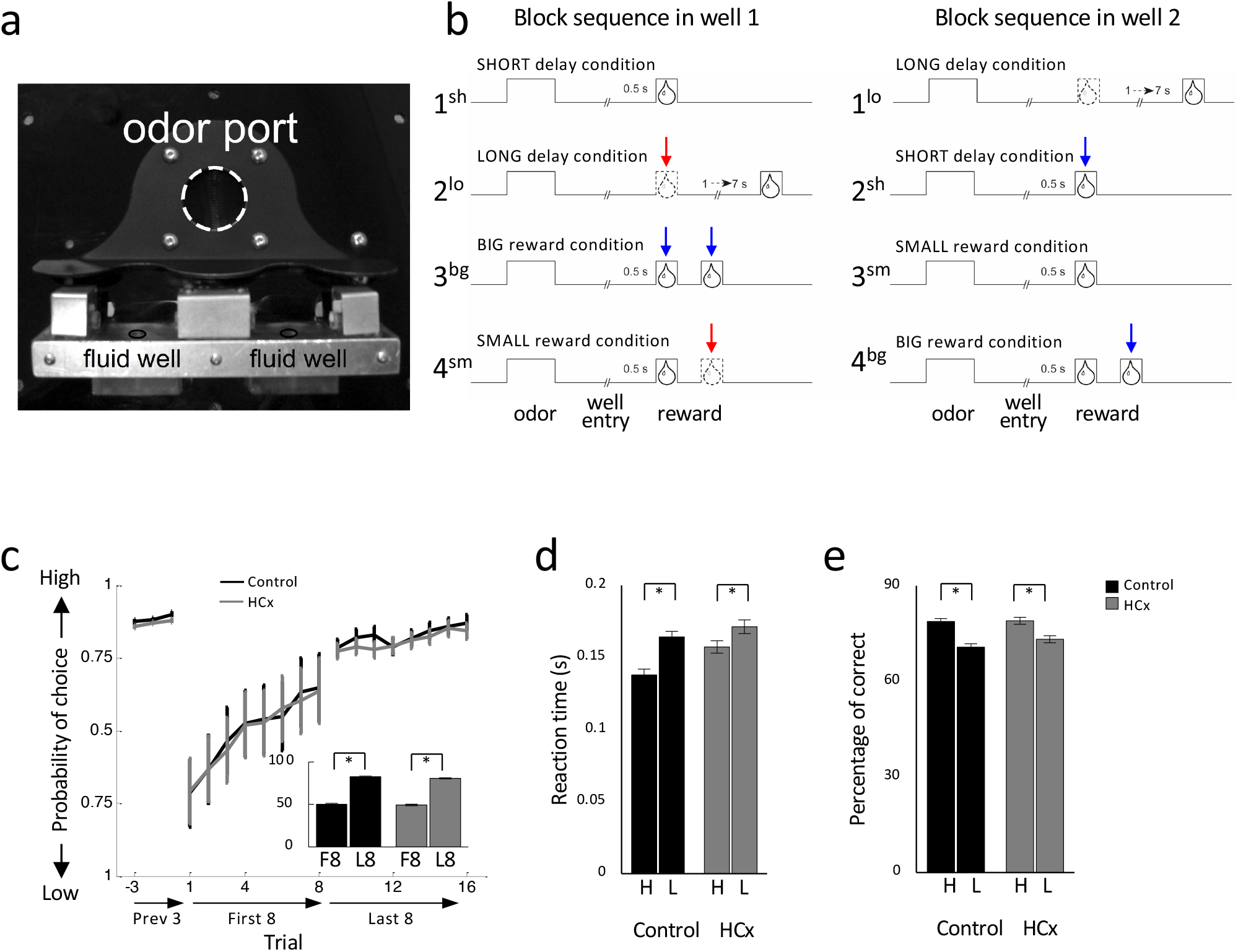
Task design and behavior. **(a)** Picture of apparatus used in the task, showing the odor port (∼2.5 cm diameter) and two fluid wells. **(b)** Deflections indicate the time course of stimuli (odor and reward) presented to the animal on each trial. Dashed lines show when a reward was omitted, and solid lines show when reward was delivered. At the start of each recording session, one well was randomly designated to deliver a single drop of reward after a short delay (0.5 sec). In the other well, one drop of reward was delivered after a long delay (a 1-7 sec). In the second block, these contingencies was switched. In third and forth blocks, delay was held constant while the number of the rewards delivered was manipulated; one well was designated as big reward in which a second bolus of the reward was delivered (big reward) and a single bolus of reward was delivered in the other well (small reward). Blue arrows, unexpected reward delivery; Red arrows, unexpected reward omission. Choice behavior in last 3 trials before the block switch, first 8 and last 8 trials after the block switch. Bar graphs indicate average percentage of choice for high valued reward in first 8 and last 8 trials after block switch. In both groups, rats chose high valued well more often on later trials than earlier trials (F’s < 861.8, p’s < 0.01). Reaction time in response to high and low valued reward on last 10 forced trials. Both groups showed faster reaction time when the high valued reward was at stake (Control, F_1,127_ = 57.9, p<0.01; HCx, F_1,160_ = 25.8, p<0.01). **(e)** percentage of correct in response to high and low valued reward on last 10 forced trials. Both groups showed higher accuracy when the high valued reward was at stake (Control, F_1,127_ = 67.0, p<0.01; HCx, F_1,160_ = 42.7, p<0.01).

Control rats changed their choice behavior across blocks in response to the changing rewards, choosing the higher value reward more often on free-choice trials (ANOVA, F_1,127_ = 575.1, p < 0.01, Fig. 1c) and responding more quickly (ANOVA, F_1,127_ = 57.9, p < 0.01, Fig. 1d) and accurately (ANOVA, F_1,127_ = 67.0, p < 0.01, Fig. 1e) on forced-choice trials when the earlier or larger reward was at stake. Rats in the HCx group also showed similar effects of value on behavior (percent choice, ANOVA, F_1,160_ = 961.8, p < 0.01, Fig. 1c; percent correct, F_1,160_ = 42.7, p < 0.01, Fig. 1e; reaction time, F_1,160_ = 25.8, p < 0.01, Fig. 1d). Inclusion of group as a factor revealed no effects of lesions on free choice trials (F’s < 1.87, p’s > 0.10), however there were significant interactions between group x value on forced-choice trials (F’s > 4.54, p’s < 0.05).

We identified putative dopamine neurons by means of a waveform analysis like that used to identity dopamine neurons in primates (Bromberg-Martin et al., 2010; Fiorillo et al., 2008; Hollerman and Schultz, 1998; Kobayashi and Schultz, 2008; Matsumoto and Hikosaka, 2009; Mirenowicz and Schultz, 1994; Morris et al., 2006; Waelti et al., 2001). This analysis isolates neurons in rat VTA whose firing is sensitive to intravenous infusion off apomorphine or quinpirole (Jo et al., 2013; Roesch et al., 2007) and which are selectively eliminated by infusion of a Casp3 neurotoxin (AAV1-Flex-TaCasp3-TEVp) into VTA of TH-Cre transgenic rats (Takahashi et al., 2017).

This approach identified as putatively dopaminergic 72 of 390 and 117 of 510 neurons recorded from VTA in control and HCx rats, respectively (Fig. S2a). These proportions did not differ between groups (Chi-square = 2.67, p > 0.10) nor were there any effects of lesions on the waveform characteristics of these neurons (Fig. S2b, t-test; Ps > 0.20). Of these, 44 neurons in control and 66 in HCx rats increased firing in response to reward (Fig. S2c, t-test, p < 0.05, compared with a 400 ms baseline taken during the inter-trial interval before trial onset). Average baseline activity was similar between the two groups for these neurons (F_1,108_ = 1.05, p > 0.10) as well as for the non-responsive dopamine neurons (F_1,77_ = 0.03, p > 0.10) and the neurons that were classified as non-dopaminergic (F_1,709_ = 1.45, p > 0.10).

### Dopamine neurons signal prediction errors in response to manipulations of reward in controls

Prediction error signaling was readily observed in response to changes in reward in dopamine neurons recorded in control rats. The activity of these neurons was elevated in response to delivery of an unexpected reward and suppressed in response to an omission of expected reward (Fig.2a). To quantify these changes in firing, we computed difference scores for each neuron by comparing the average firing at the beginning versus the end of the blocks at the time of reward delivery or omission. The distributions of these scores were shifted above zero when unexpected reward was delivered (left in Fig. 2c) and below zero when expected reward was omitted (right in Fig. 2c), reflecting that changes in firing (elevation or suppression) were maximal at the start of the block and then diminished with learning of the new contingencies as the block proceeded (Fig 2d).

**Figure 2:**
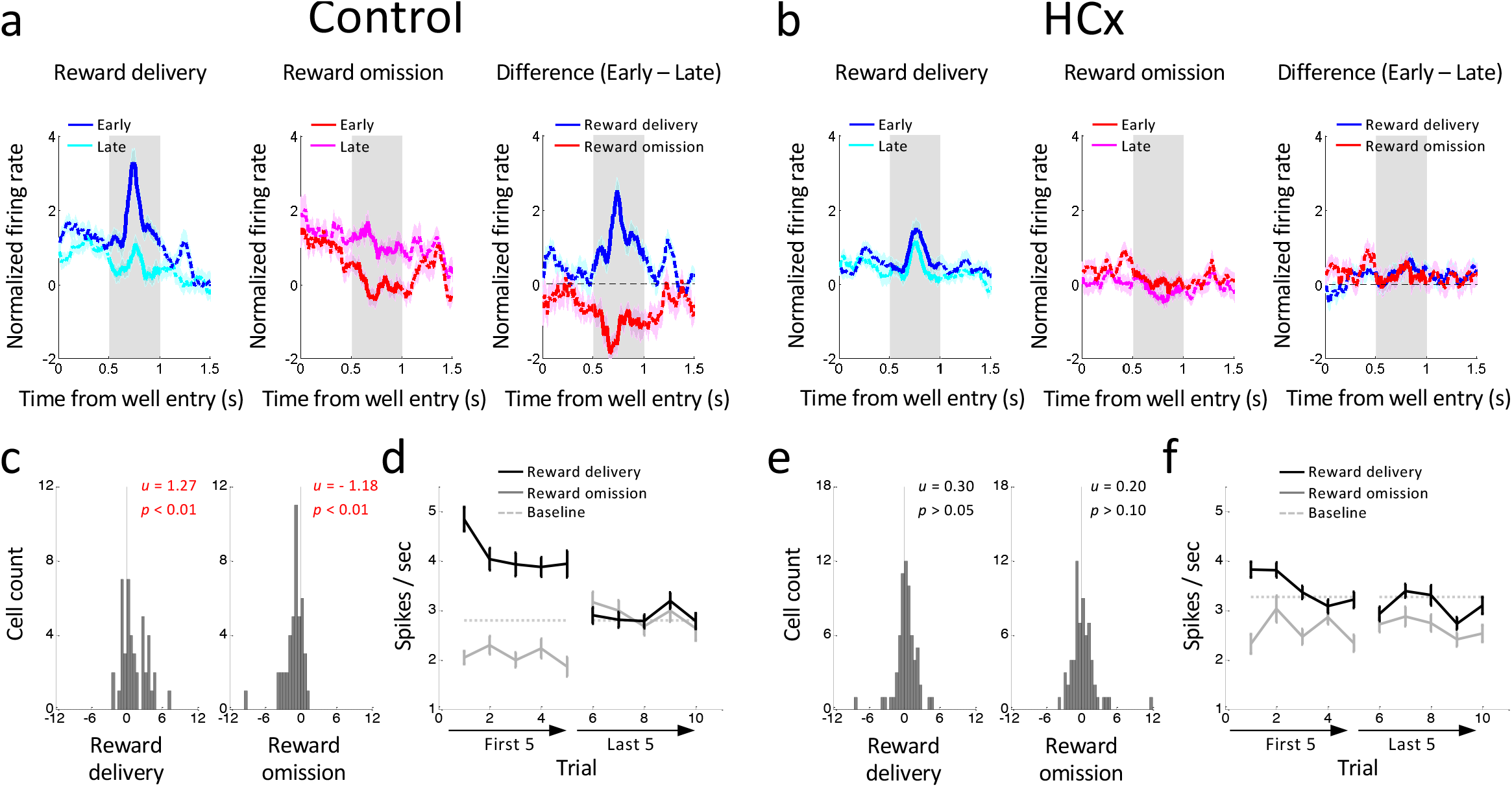
Changes in reward-evoked activity of reward-responsive dopamine neurons to changes in reward value. **(a and b)** Population responses of reward-responsive dopamine neurons in Control (a) and HCx (b) groups. Left panels show changes in firing to reward delivery on the first (dark-blue) and last (light-blue) trials. Middle panels show changes in firing to reward omission on the first (dark-red) and last (light-red) trials. Right panels show the difference in firing between first and last trials in response to reward delivery (blue) and omission (red). **(c and e)** Distributions of difference scores comparing firing to unexpected reward delivery (left) and omission (right) in the early and late trials in control (c) and HCx (e) groups. The numbers in each panela indicate results of Wilcoxon singed-rank test (p) and the average difference score (u). **(d and f)** Average firing in response to reward delivery (black) and omission (gray) in the first 5 and last 5 trials in control (d) and HCx (f) groups. ANOVA (Reward x Early/Late x Trial) revealed significant main effects of Reward (Control, F_1,43_ = 13.0, p<0.01; HCx, F_1,65_ = 23.0, p<0.01) and Trial (Control, F_1,43_ = 2.57, p<0.05 ; HCx, F_1,65_ = 5.09, p<0.01) in both control and HCx. and a significant interaction of Reward x Early/Late in control (F_4,172_ = 44.1, p<0.01), but not in HCx (F_4,260_ = 1.95, p>0.10). A step down in each plot revealed a significant main effect of Early/Late in reward delivery in both groups (control, F_1,43_ = 27.0, p<0.01; HCx, F_1,65_ = 4.85, p<0.05) and reward omission in control (F_1,43_ = 14.8, p<0.01), but not in HCx (F_1,65_ = 0.08, p>0.10). Dashed lines indicate baseline firing. Error bars, S.E.M.

### Ipsilateral hippocampal lesions disrupt error signaling to reward manipulations by dopamine neurons

HC lesions had a marked effect on the changes in the firing of dopamine neurons caused by changes in reward. In particular, dopamine neurons recorded in rats with ipsilateral HC lesions did not increase firing when reward was delivered unexpectedly (left in Fig. 2b) nor did they suppress firing when an expected reward was omitted (middle in Fig. 2b). These effects were again quantified by analyzing the difference scores between early versus late trials in relevant blocks. The difference scores were not different from zero when an unexpected reward was delivered (Fig. 2e left) or an expected reward was omitted (Fig. 2e right), reflecting the relatively flat firing in early and late trials of each type (Fig. 2f).

Consistent with these apparent differences, direct comparison data from control and HCx-lesioned rats (ANOVA, group x reward/omission x early/late x trial, Fig. 2d versus Fig. 2f) revealed significant group interactions (group x reward/omission x early/late; F_4,432_ = 13.4, p < 0.01), and the distributions of the difference scores comparing firing changes in reward delivery or omission (histograms, Fig. 2c versus 2e) were significantly different between the group (Wilcoxon rank-sum test; ps < 0.05). Thus dopamine neurons recorded in rats with ipsilateral HC lesions failed to show normal bidirectional changes in firing – presumed to be reward prediction errors -in response to manipulations of reward.

### Ipsilateral hippocampal lesions had no effects on cue-firing in VTA dopamine neurons

The activity of dopamine neurons in control rats also differed during sampling of the odor cues according to the expected value of the cue in the different blocks. On forced-choice trials, these neurons exhibited higher firing during the presentation of the high-valued cue than during presentation of the low valued cue, a difference that reversed in each block early in learning (Fig. 3a). To quantify this, we computed the difference scores comparing each neuron’s firing to the high-versus low-value cues in early versus late trials. Distribution of these scores was shifted significantly above zero (Fig. 3b) in controls, indicating an increase/decrease in activity to the high/low value cues across trials, respectively.

**Figure 3:**
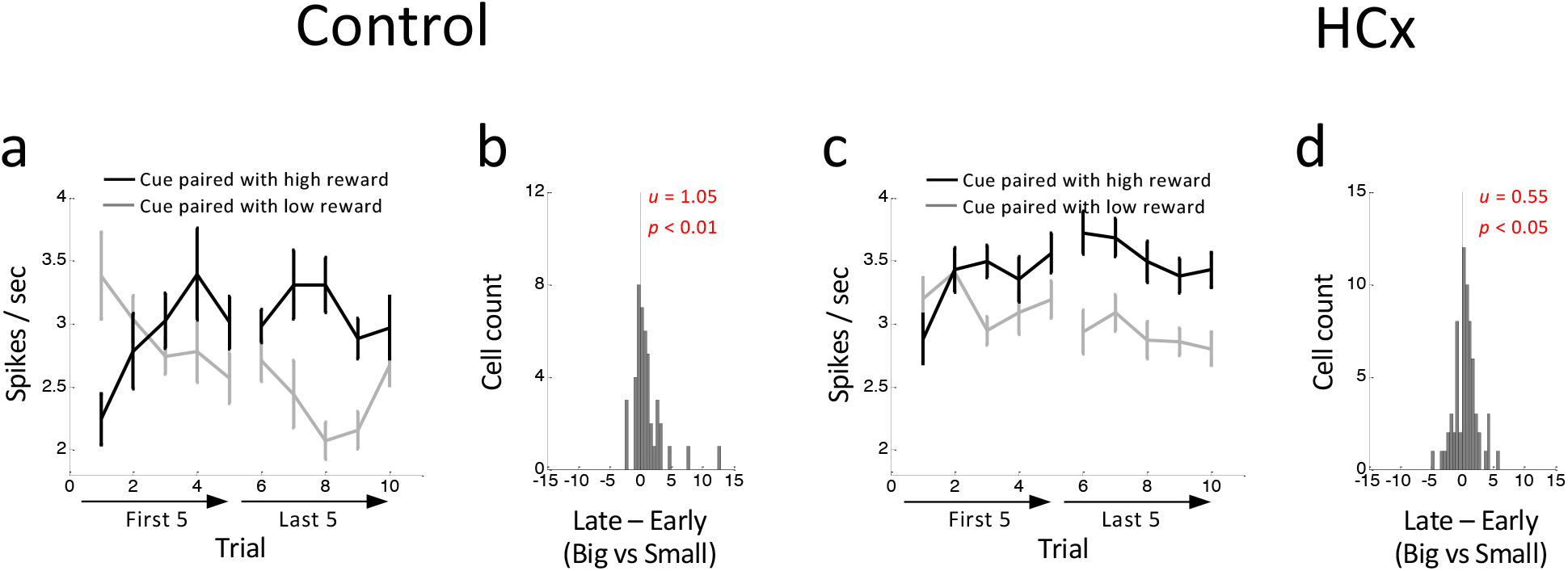
Changes in reward-evoked activity of reward-responsive dopamine neurons to reward predictive odor cues. **(a and b)** Average firing in response to high-(black) and low-valued (gray) cues in the first 5 and last 5 trials in control (a) and HCx (b) groups. ANOVA (group x value x early/late x trial) revealed a significant main effect of value (F_1,108_ = 19.8, p<0.01) and significant interactions of value x early/late (F_1,108_ = 12.7, p<0.01), value x trial (F_4,432_ = 3.87, p<0.01), and value x early/late x trial (F_4,432_ = 2.64, p<0.05). Error bars, S.E.M. **(b and d)** Distributions of difference scores between high- and low-valued cues in early and late trials in control (b) and HCx (d) groups. The numbers in each panela indicate results of Wilcoxon singed-rank test (p) and the average difference score (u).

Somewhat surprisingly, similar effects were also evident in dopamine neurons recorded in rats with ipsilateral HC lesions (Fig. 3c-d). A direct comparison of the data between control and lesioned groups (group x value x early/late x trial) revealed a significant main effect of value (F_1,108_ = 19.8, p < 0.01) and significant interactions between value x early/late (F_1,108_ = 12.7, p < 0.01), value x trial (F_4,432_ = 3.87, p < 0.01) and value x early/late x trial (F_4,432_ = 2.64, p < 0.05). However, there were no significant main effects nor interactions involving group (F’s < 1.5, p’s > 0.10). Thus dopamine neurons recorded from rats with ipsilateral HC lesions showed normal changes in firing in response to presentation of the differently-valued cues.

### Hippocampal lesions disrupt hierarchical segregation of states available in different blocks

The neural results showed that HC was necessary for normal error signaling by VTA dopamine neurons; dopamine neurons recorded in rats with ipsilateral HC lesions failed to signal prediction errors to changes in reward, while at the same time showing roughly normal error signals to the presentation of the predictive cues in our task. To better understand what hippocampus might contribute to cause this pattern of results, we developed temporal-difference reinforcement learning models to describe the task and then monitored the error output of the model caused by changes in several parameters in an attempt to recreate these neural findings.

We modeled a learning agent that represents the behavioral task as serial transitions between hidden states within a partially-observable semi-Markov process (Daw et al., 2006). In this model, states were each associated with a finite duration, captured by a distribution over possible “dwell” times within a given state. States were also probabilistically associated with unique observations made during the task (for example, the onset of a cue, or entry into one of the reward wells), such that these events mark the start of the dwell time within a state. While observations unambiguously signal transitions between states, transitions could also happen without observations. As in other partially-observable settings, a belief state—that is, a probability distribution over state occupancy—tracks the agent’s understanding of their current position within the various possible states of the task (Starkweather et al., 2018). In order to estimate the current state and learn about its value (i.e. the expected future reward), the agent had to combine task observations (i.e. the odor presented, number of rewards), the duration of delays since the last observation (i.e. whether short or long), and their current knowledge of the underlying map of the task (i.e. transitions between states and their associated dwell time distributions). If the agent inferred that a state transition had occurred, then a prediction error was calculated based on the states’ current estimated value, the reward feedback, and the probability of that state transition having just occurred. The value of the previous state was then updated through a temporal-difference learning rule, and, in parallel, the dwell time distribution was updated to reflect the time spent in that state.

Using this basic architecture, we developed two different models to explain the effects of HC lesions. The first model was simpler, reflecting only the states and dwell time distributions the rats experienced when performing the task (Fig. 4a-b, model 1). In this model, the agent learned the value and dwell times of each state through experience, to minimize the deviation between expectations and observations. This resulted in a pattern of reward prediction errors at the time of reward delivery and cue sampling very similar to that observed in dopamine neurons recorded from control rats (Fig. 4c).

**Figure 4:**
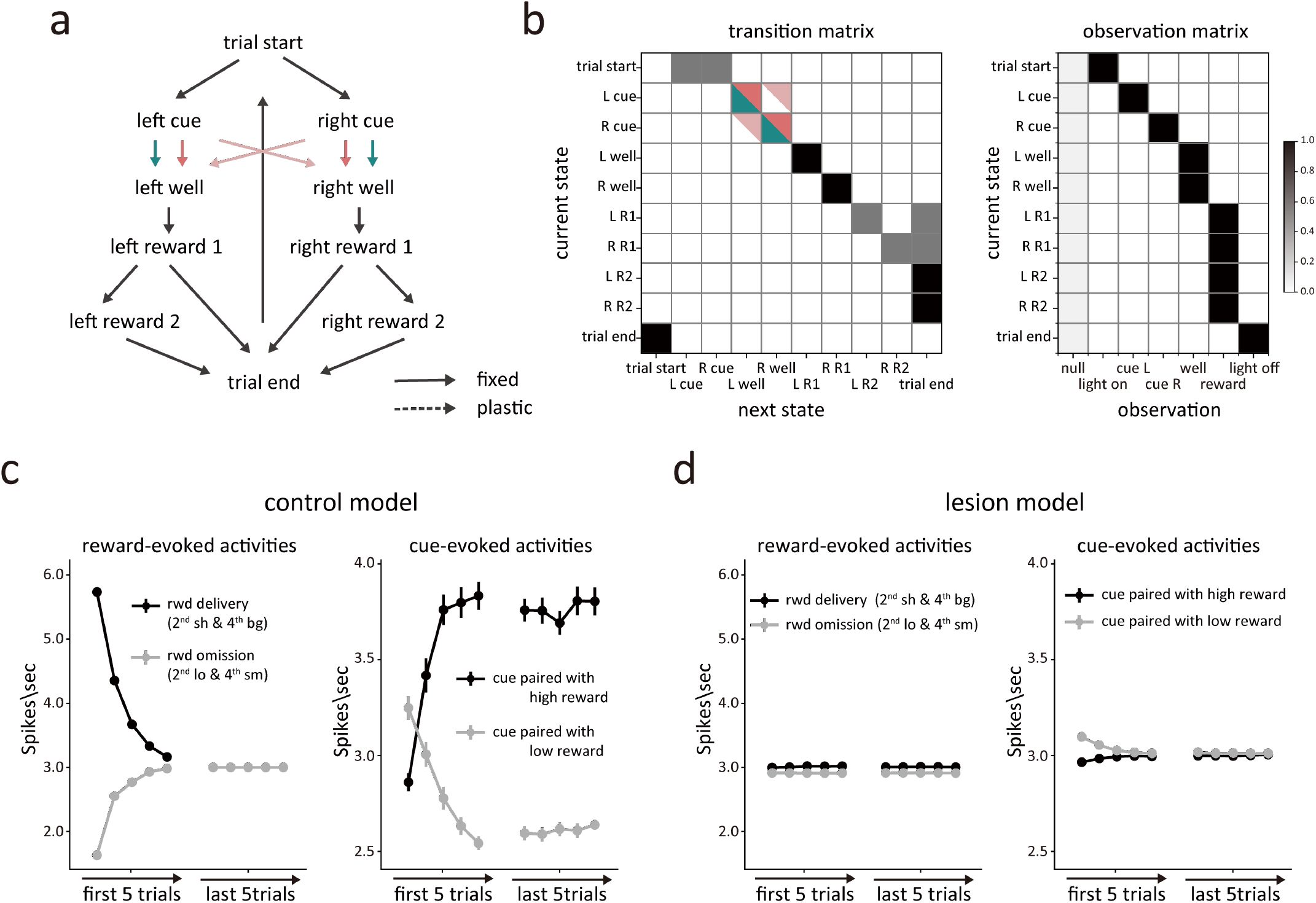
Modeling the effect of hippocampal lesions as a blurring of transitions between trials. **(a)** Conventional state space representation of the task. Possible transitions between states are depicted by arrows. Green arrows represent transitions available to the control model, while pink arrows represent transitions available to the lesion model. **(b)** The transition matrix (left panel) shows the probability of each successor state given each state, and the observation matrix (right panel) shows the probability of each observation given each state. The darker color indicate higher probabilities. Green and pink indicate the transitions available to the control and lesioned models, respectively. The characteristic observation is emitted with p = 0.95. States also emit a null (empty) observation (p = 0.05) or any of the other five possible observations (with p = 1e-4 each). The observation probabilities for each state were normalized by dividing their sum. **(c)** Simulated average prediction errors in the control model during the 2^*nd*^ and 4^*th*^ blocks. In the left panel, the black and grey lines represent the prediction error in response to reward delivery and reward omission, respectively. In the right panel, the dark and light line represents the prediction error in response to odor cue paired with high and low reward, respectively. **(d)** The same format as Fig. 4c, but for the lesion model.

To model the effects of HCx in this setting, we elected to blur the ability of the model to maintain internal information about the transition probabilities between the odor cues and the correct well states (pink arrows in Fig. 4a). This reflected the hypothesis that HC lesions would disrupt the maintenance of hidden information during the period after a response had been made, while the rats were waiting for reward, an effect like that caused by lesions of OFC in this task (Takahashi *et al*., 2011). In the context of the model, this change caused the agent to rely more heavily on external observations and dwell time than on the transition matrix when estimating the current state. As a result, the agent was more likely to respond to unexpected external input regarding events or timing – like the unexpected appearance of reward (Fig. 1b, 2^*sh*^ or 4^*bg*^) or its unexpected delay or omission (Fig. 1b, 2^*lo*^ or 4^*sm*^) -by changing its estimate of the current state to the opposite well (Fig. 4b, light pink shading). This reevaluation resulted in the loss of prediction error signaling by the agent in these blocks, since it essentially brought their belief state and its predictions into alignment with external events (Fig. 4c, left panel). This result aligns well with main effects of lesions on dopamine neuron firing in the task – specifically the apparent loss of responsivity to unexpected reward and reward omission (Fig. 2).

However, this model predicted something very different at the transition between blocks 2 and 3. Because block 3 is the first block in the session that involves two drops of reward, there is no internal state at the end of block 2 whose value estimate can predict the delivery of the second reward (Fig 1b., 3^*bg*^). As a result, the lesioned model produced a strong positive prediction error to the second reward on these trials (Fig 6a). This effect was not evident in the activity of dopamine neurons recorded in the HCx rats (Fig. 6a). Further the lesioned model also failed to produce the apparently normal error responses to the odor cues observed in the dopamine neurons recorded in HCx rats (Fig 4d, right panel).

Given the poor match between the output of the simplified lesion model and the data, we developed a second model, based on the same semi-Markov process but a more complex and more realistic state space that recognized the extensive training history of the rats on the task by creating separate clusters of states for each of the different blocks, reflecting the unique reward contingencies of each block. This resulted in a multi-level or hierarchical state space in which lower-level states described the underlying process within individual trials and upper-level states controlled the probabilities of transitions to different blocks based on recent reward history (Fig. 5a). As with the simpler model, this model did a good job reproducing the pattern of reward prediction errors evident at the time of reward delivery and cue sampling in dopamine neurons recorded from control rats (Fig. 5c).

**Figure 5:**
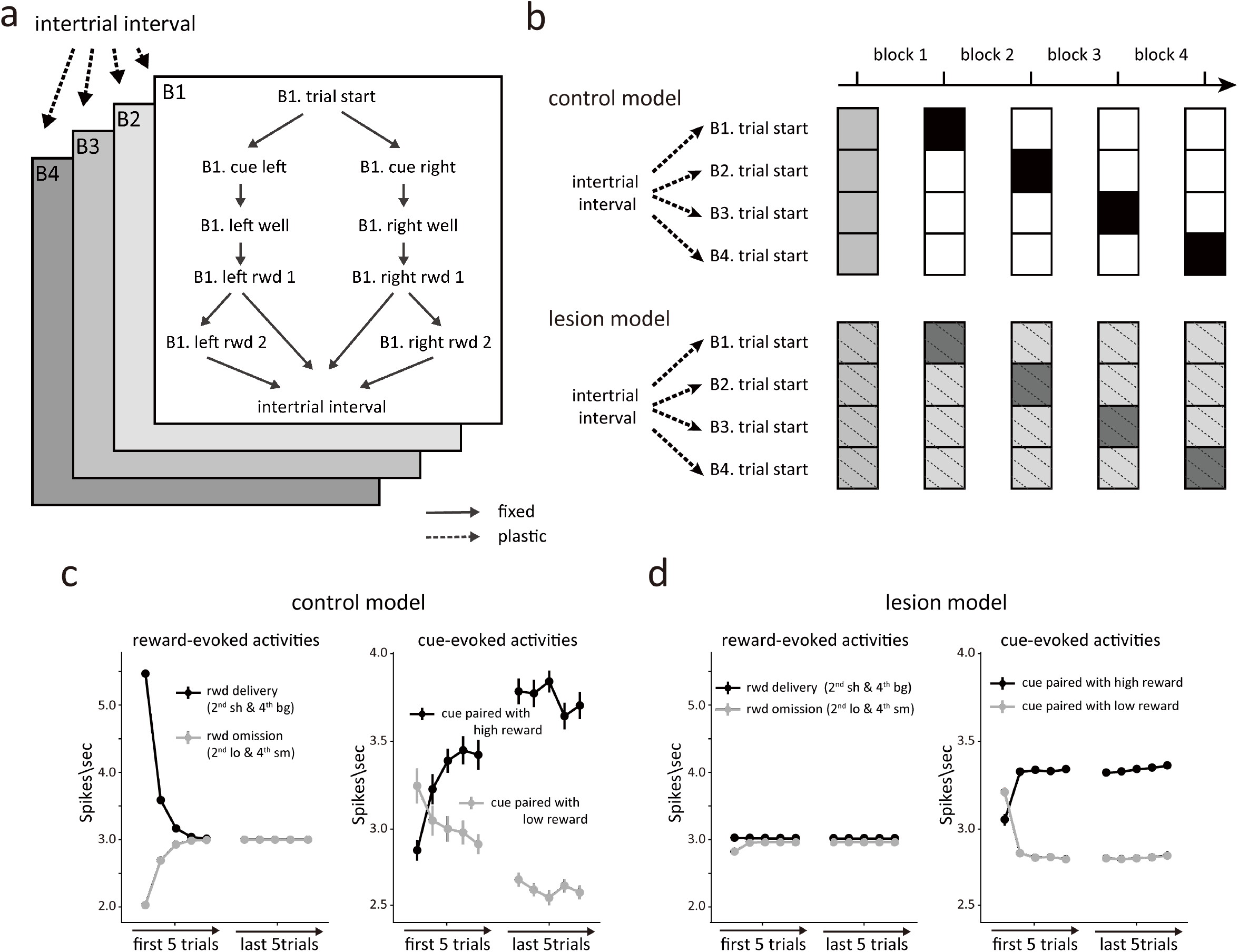
Modeling the effect of hippocampal lesions as a blurring of transitions between blocks. **(a)** Multi-level or hierarchical state space representation of the task. The upper-level contains the transition from intertrial interval to each block, with the trial start state in each block leading to the lower-level states, describing the state space of individual trials. Available transitions between states are marked by arrows. Dashed arrows indicated plastic transitions, whose probabilities are updated during learning. Solid arrows are transitions with fixed probabilities. **(b)** The transition probabilities in the upper-level of the task space are updated according to reward history. The control model learns the transition probabilities perfectly and with low uncertainty (top panel), while the lesioned model has greater residual uncertainty (bottom panel). The darker color indicate higher probabilities. **(c)** Simulated average prediction errors in the control model during the 2^*nd*^ and 4^*th*^ blocks. In the left panel, the dark and light line represents the prediction error in response to reward delivery and reward omission, respectively. In the right panel, the black and grey lines represent the prediction error in response to odor cue paired with high and low reward, respectively. **(d)** The same format as Fig. 5c, but for the lesion model.

**Figure 6:**
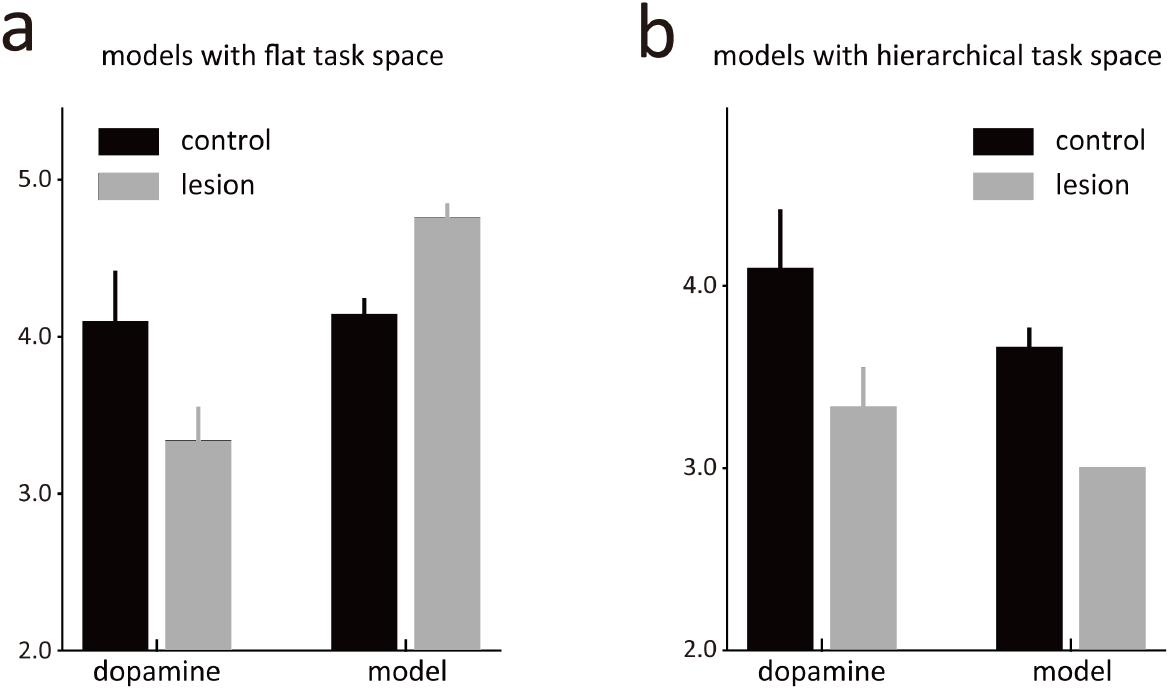
The model 2 with a hierarchical task space provides a better explanation for dopamine neuron activities. **(a)** Comparison of activities evoked by the 2nd drop of reward in the first 5 trials of the 3^*rd*^ block in both animals and the model 1 with a flat task space. The control animals’ dopamine neurons exhibit a higher firing rate compared to the HCx animals. In contrast, the model 1 with a flat state space shows the opposite pattern. **(b)** Same format as the Fig. 6a, but for the model 2 with a hierarchical task space. The firing pattern in the model 2 is consistent with dopamine neuron activities.

Using this more complex state space, we again modeled the effect of hippocampal lesions as a blurring of transitions, but this time between the upper-level states describing the trial blocks (Fig. 5b). This reflected the hypothesis that hippocampus might differ from OFC in that it would be necessary for maintaining hidden information only when events were discontinuous or separated by longer timescales. To implement this, we introduced larger uncertainty in the transition probabilities between the upper-level states (Fig. 5b), which were updated based on recent reward history in the lesioned model. Though counterintuitive at first glance, the imprecise transition probabilities caused the lesioned model to again be more heavily influenced by external observations about events and dwell times, in this case affecting its estimate about the current block. As a result, the lesioned model was again able to quickly adapt to changes in reward number or timing by altering its estimated state rather than adjusting through error signaling, but unlike the prior model, this ability now included the third block where two rewards were introduced (Fig. 6b).

This model also captured the relative preservation of error signals in response to the odor cues observed in the HCx rats (Fig 3). This was possible because the lesioned model updated the transition probabilities based on reward history, leading to an update in the estimated belief state behind the odor cues after the initial trial in a block and resulting in changes in cue-evoked activity (Fig. 5d, right panel). Interestingly, the lesioned model also exhibited clear differences in the cue-evoked errors that arguably mirrored minor but apparent differences in the preserved responses in HCx rats; in particular, while not significant, the cue-evoked response in these rats seemed weaker and to change more completely after the initial trial than the response in controls (Fig. 3). Overall, this second model based on blurring of upper-level internal information about the states available in different blocks effectively captured the characteristics of dopamine neurons’ firing, suggesting that HC-lesioned animals may be deficient in maintaining and updating higher-order state representations that capture longer timescale task structure within a cognitive map.

## Discussion

In the current study, we utilized the well-characterized dopaminergic reward prediction error as a tool to investigate the critical role of the HC in the development of cognitive or task maps for guiding learning from direct experience of rewards, repeating an approach used previously to characterize the role of OFC in cognitive mapping (Takahashi *et al*., 2011; Wilson *et al*., 2014). We recorded the activity of midbrain dopamine neurons during the induction of prediction errors in an odor-based choice task and compared activity in controls with the activity in rats with ipsilateral HC lesions. Combined with computational modeling, this allowed us to gain insight into the specific effects of HC removal on the structure of the internal task representation being used for learning and compare them to the effects of OFC lesions. The results show that HC, like OFC, contributes to representation of the appropriate state structure in this task, at least with regard to the information available to midbrain dopamine neurons, and appears to be especially important for properly representing information that is hidden or at least partially observable. However, unlike OFC, this contribution is not required for properly segregating states within the trial but instead becomes important at longer timescales, across trials, where it is critical to organizing information about the context or current block.

In evaluating these results, it is worth noting that the models used in the current study are largely the same as the model used previously to understand the effects of OFC lesions in this task (Takahashi *et al*., 2011), aside from the addition of a semi-Markov process added to allow the model to learn about reward timing and selectively disrupted by ventral striatal lesions (Takahashi et al., 2016). The current models inherited the basic task architecture from the prior work on OFC, although the second model was extended with a HC-dependent hierarchical element to accommodate separate representations of the states available in different blocks of trials. Critically, by modifying either the transition matrix or dwell time distribution in the basic model used here, we can replicate the changes in dopamine neuron firing observed in prior work after OFC or ventral striatal lesions (see Figure S3). Thus, the results here are not due to idiosyncratic differences between the models and do not alter conclusions drawn in those prior studies.

The current findings are significant both for contrasting the roles of HC and OFC in cognitive mapping as well as for revealing the complexity of information afferent to the midbrain dopamine neurons. With regard to HC and OFC function, the results are consistent with the idea that both areas are critical for constructing maps of complex task spaces, but within the constraints of our task they point to different contributions. The OFC was particularly important for segregating states – in this case the left and right wells – significant to reward (i.e., one direction is rewarded, and one is not on forced-choice trials, and one is always better on free choice) and negligible external differences (both wells are identical). This function is consistent with evidence in other settings that the OFC is particularly critical for representing latent, hidden, or partially observable information of potential behavioral relevance (Costa et al., 2023; Schuck et al., 2016; Wilson *et al*., 2014). By contrast, the HC was not necessary for maintaining this information, however it played a key role in properly representing the current context or trial block. Information identifying the different trial blocks is also hidden or partially observable (i.e. there are no unique events that identify different blocks in the ITI or even during the trial until actual reward), however properly maintaining information about trial block has a strong temporal component requiring memory, thus the dependence on HC is clearly consistent with the historical role of HC in episodic memory and contextual information (Duvelle et al., 2022; Farovik et al., 2015; Smith and Bulkin, 2014), as well as more recent evidence that the HC encodes time as well as parsing statistical regularities that are critical to constructing hierarchical task spaces (Schapiro et al., 2016; Stachenfeld et al., 2017). Which one of these features is critical for the involvement of the HC in our task is difficult to say, however it is tempting to suggest it is the hierarchical, reward-orthogonal organization (Farovik *et al*., 2015). In complex environments with an overwhelming number of states, agents – humans or rats – are theorized to organize the cognitive map and plan actions hierarchically for efficient behavior (Botvinick, 2012). Yet most behavioral tasks used in behavioral neuroscience are not obviously hierarchical nor are the results analyzed to consider possible hierarchical solutions. This makes it difficult to ascertain whether the HC encodes states at all orders equally or exhibits greater encoding of upper-level states. Although our findings indicate that the HC lesion disrupts the clear delineation of transitions between upper-level states, we do not intend to suggest that the HC is not involved in encoding other lower-level states or their transitions. Rather, other brain regions such as the OFC may perform similar functions allowing compensation.

With regard to dopamine function, these results highlight the potential complexity of sources and information available to this powerful teaching system. VTA dopamine neurons receive information from numerous brain areas, both directly and indirectly (Watabe-Uchida et al., 2012). Although these various inputs may be redundant when behavioral demands are low (Watabe-Uchida *et al*., 2012), as the dopamine firing pattern can still be well reconstructed even when input from a single area is removed *in silico*, the current results add to a growing body of data indicating that redundant coding may be limited to these simple situations, in which hidden structure is not important in determining the internal cognitive map according to which predicted values are calculated. In more complex situations, the unique contributions of different brain regions would be expected to – and clearly do – start to emerge. This points to the importance of studying the specific contributions of brain regions in behavioral contexts sufficiently complex to test their unique contributions.

Having access to rich sources of information regarding the task structure is also important because, though it is much ignored, the complexity of the internal state representation can have a dramatic influence on what sorts of behaviors can be supported by both the actual dopamine system and by the temporal difference learning rule which has been mapped onto it (Akam et al., 2015). While much work in the field focuses on showing that current accounts either can or cannot explain certain findings, it is perhaps worthwhile considering whether much of the uncertainty or contradictory results could be explained by a better understanding of how individual animal subjects represent the state space of a task, along with the biases and idiosyncrasies that are native to volitional behavior. For instance, it has been suggested that contingency degradation provides an insurmountable challenge to the temporal difference reinforcement learning algorithms (Dezfouli and Balleine, 2012), since an agent armed with a very simple state space that considers only the cue will learn the value of a cue based on its contiguity with reward, without impact of non-contingent reward delivery during periods when the cue is not present (Garr et al., 2023; Jeong et al., 2022). However, if this agent has access to a state space in which the context is considered and allowed to be a target and compete for learning – including when the cue is present – then a temporal difference rule will in principle reproduce the known effects of contingency degradation (i.e. delivery of rewards during non-cue periods) on behavior and dopamine release to a degraded cue.

Indeed, the ability of the dopamine system to support learning and behavior in the real world, which is fantastically complicated compared to even our task, likely depends entirely on the complexity and detail of the input the system receives. A teaching signal is only as good as the information it receives about the world. This is increasingly being recognized by the incorporation of channels, features, and bases into temporal difference learning models (Lee et al., 2022; Millidge et al., 2023; Takahashi et al., 2023) and in research positing that dopamine neurons are influenced by internal information, inference, and dynamically evolving beliefs (Hennig et al., 2023; Lak et al., 2017; Nomoto et al., 2010; Papageorgiou et al., 2016; Sadacca et al., 2016; Starkweather et al., 2017; Starkweather *et al*., 2018; Wassum et al., 2011). Yet these findings are likely only the tip of the iceberg; the results here and in the related studies show experimentally that information from even high-level association cortices is utilized by these neurons.

## Methods

### Lead Contact

Geoffrey Schoenbaum.

### Data and Code Availability

The dataset and all scripts used in this study will be made available in an appropriate data archive upon publication.

### Experimental Model and Subject Details

Data from fourteen male Long-Evans rats (Charles River Labs, Wilmington, MA) contributed to this study; this does not include two rats that expired post-operatively during recording whose data were not used. Rats were tested at the NIDA-IRP in accordance with NIH guidelines.

### Method Details

To allow direct comparisons to be made to prior work, all equipment and procedures used were substantially the same to those used in a prior study in which dopamine neurons were recorded in rats with ipsilateral lesions of the OFC (Takahashi *et al*., 2011). In particular, the approach to recording dopamine neurons, the electrode design, recording systems, general task, specific task and training procedures, strain, age and sex of rat used were all identical to the prior study.

### Stereotaxic Surgery

All surgical procedures adhered to guidelines for aseptic technique. For electrode implantation, a drivable bundle of eight 25-um diameter formvar insulated nichrome wires (A-M systems, Carlsborg, WA) chronically implanted dorsal to VTA in the left or right hemisphere at 5.3 mm posterior to bregma, 0.7 mm laterally, and 7.5 mm ventral to the brain surface at an angle of 5° toward the midline from vertical. Some rats (n = 9) also received neurotoxic lesion of ipsilateral hippocampus by the infusion of NMDA (20 mg/ml) at seven sites in each hemisphere (see Fig. S1 for the surgical coordinates). Controls (n = 5) received sham lesions in which burr holes were drilled and the pipette tip lowered into the brain but no solution delivered. Cephalexin (15 mg/kg p.o.) was administered twice daily for two weeks post-operatively.

### Histology

All rats were perfused at the end of the experiment with phosphate-buffered saline (PBS) followed by 4% paraformaldehyde (Santa Cruz Biotechnology Inc., CA). Brains were cut in 40 μm sections and stained with thionin.

### Odor-guided choice task

Recording was conducted in aluminum chambers approximately 18” on each side with sloping walls narrowing to an area of 12” x 12” at the bottom. A central odor port was located above two fluid wells (Fig. 1a). Two lights were located above the panel. The odor port was connected to an air flow dilution olfactometer to allow the rapid delivery of olfactory cues. Odors were chosen from compounds obtained from International Flavors and Fragrances (New York, NY). Trials were signaled by illumination of the panel lights inside the box. When these lights were on, nosepoke into the odor port resulted in delivery of the odor cue to a small hemicylinder located behind this opening. One of three different odors was delivered to the port on each trial, in a pseudorandom order. At odor offset, the rat had 3 seconds to make a response at one of the two fluid wells. One odor instructed the rat to go to the left to get reward, a second odor instructed the rat to go to the right to get reward, and a third odor indicated that the rat could obtain reward at either well. Odors were presented in a pseudorandom sequence such that the free-choice odor was presented on 7/20 trials and the left/right odors were presented in equal numbers. In addition, the same odor could be presented on no more than 3 consecutive trials. Once the rats were shaped to perform this basic task, we introduced blocks in which we independently manipulated the size of the reward or delay preceding reward delivery (Fig. 1b). For recording, one well was randomly designated as short and the other long at the start of the session (Fig. 1b, 1^*sh*^ and 1^*lo*^). In the second block of trials, these contingencies were switched (Fig. 1b, 2^*sh*^, 2^*lo*^). The length of the delay under long conditions followed an algorithm in which the side designated as long started off as 1 s and increased by 1 s every time that side was chosen until it became 3 s. if the rat continued to choose that side, the length of the delay increased by 1 s up to a maximum of 7 s. If the rat chose the side designated as long less than 8 out of the last 10 choice trials, then the delay was reduced by 1 s to a minimum of 3 s. The reward delay for long forced-choice trials was yoked to the delay in free-choice trials during these blocks. In the third and fourth blocks we held the delay preceding reward constant while manipulating the number of reward (Fig. 1b, 3^*bn*^, 3^*sm*^, 4^*bg*^ and 4^*sm*^). The reward was a 0.05ml bolus of 10% sucrose solution. The reward number used in delay blocks was the same as the reward used in the small reward blocks. For big reward, an additional bolus was delivered after gaps of 500ms. The blocks were more than 60 trials long and block switches occurred when rats chose high value side more than 60% in last 10 free-choice trials.

### Single-unit recording

Wires were screened for activity daily; if no activity was detected, the rat was removed, and the electrode assembly was advanced 40 or 80 μm. Otherwise active wires were selected to be recorded, a session was conducted, and the electrode was advanced at the end of the session. Neural activity was recorded using Plexon Multichannel Acquisition Processor systems (Dallas, TX). Signals from the electrode wires were amplified 20X by an op-amp headstage (Plexon Inc, HST/8o50-G20-GR), located on the electrode array. Immediately outside the training chamber, the signals were passed through a differential pre-amplifier (Plexon Inc, PBX2/16sp-r-G50/16fp-G50), where the single unit signals were amplified 50X and filtered at 150-9000 Hz. The single unit signals were then sent to the Multichannel Acquisition Processor box, where they were further filtered at 250-8000 Hz, digitized at 40 kHz and amplified at 1-32X. Waveforms (>2.5:1 signal-to-noise) were extracted from active channels and recorded to disk by an associated workstation

### Data analysis

Units were sorted using Offline Sorter software from Plexon Inc (Dallas, TX). Sorted files were then processed and analyzed in Neuroexplorer and Matlab (Natick, MA). Dopamine neurons were identified via a waveform analysis. Briefly cluster analysis was performed based on the half time of the spike duration and the ratio comparing the amplitude of the first positive and negative waveform segments. The center and variance of each cluster was computed without data from the neuron of interest, and then that neuron was assigned to a cluster if it was within 3 s.d. of the cluster’s center. Neurons that met this criterion for more than one cluster were not classified. This process was repeated for each neuron. The putative dopamine neurons that showed increase in firing to reward compared to baseline (400ms before reward) were further classified as reward-responsive (t-test, p< 0.05). To analyzed neural activity to reward, we examined firing rate in the 400 ms beginning 100 ms after reward delivery.

### Computational models

We utilized a temporal-difference reinforcement learning (TDRL) algorithm within a partially-observable semi-Markov framework to simulate the evolution of reward prediction, and reward prediction errors, with experience on the behavioral task (Daw *et al*., 2006; Takahashi *et al*., 2016). In this model, states *s* are not observable (i.e., directly accessible to the behavioral agent), rather they are probabilistically associated with a set of observations corresponding to task events, such as the onset of the odor cue or the delivery of the reward. Each state is also probabilistically associated with a finite duration, *d*, that initiates at the time of the observation/s associated with a given state. We assumed that task events, modeled as non-empty observations, reliably signal transition to a new state, but that transitions may also occur with no corresponding event, i.e. an empty observation. Similar to other models in a partially observable setting, the conditional probabilities of each observation given a hidden state are specified by an observation function *O*, and the probability of one state following another by a transition matrix, *T*. As rats were trained extensively on the odor-guided choice task, we assumed that they had learned a cognitive model for these aspects of the task, and therefore, function *O* and transition matrix *T* were known.

In this partially observable semi-Markov TDRL model, credit assignment requires estimating the current hidden state, which we model using an inference process that tracks the probability of having just transitioned out of state *s* at time *t*, given the sequence of observations up to t+1, *β*_*s,t*_ (Daw *et al*., 2006). As state transitions occur irregularly (as opposed to on every time point), performing this inference depends critically on an estimate of the likely duration d of each state, captured by the dwell time distribution, *D* = *P*(*d*|*s*). To compute *β*_*s,t*_, we rewrited it using Bayes’ Rule:

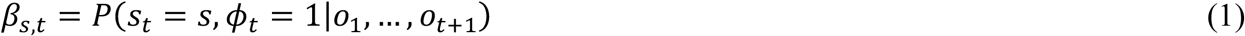

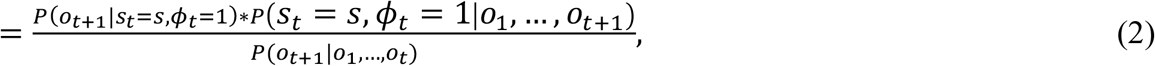

where the dummy variable, *ϕ*_*t*_, indicates whether a state transition happened at time *t*. According to the Markov property, the first term of the numerator in Eq. 2 is equal to the integration over state *s*_*t*_, i.e., 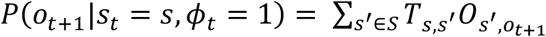. Leveraging the assumption that *β*_*s,t*_ = 1 if *o*_*t+1*_ is non-empty, the second term of the numerator, denoted *α*_*s,t*_, can be computed by integrating over all possible dwell times in state *s*, since the last non-empty observation:

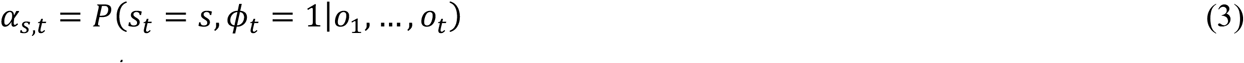

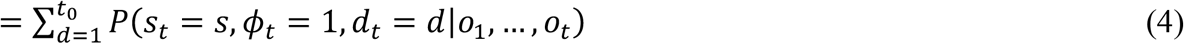

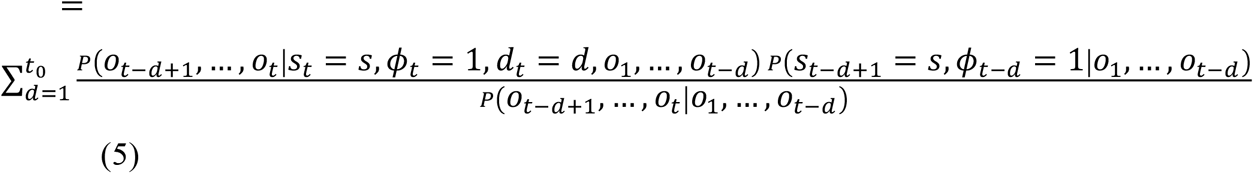

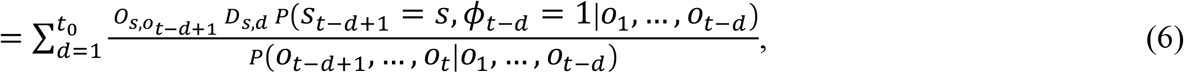

where *t*_*0*_ is the time that passed since the last non-empty observation. The last term of the numerator in Eq. 6 is the probability that the process left state *s* at time *t− d* weighted by the transition probability from *s*_*t−d*_ to *s*_*t−d+1*_, which is *α*_*s,t−d*_. Thus, *α*_*s,t*_ can be computed recursively,

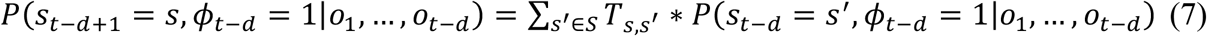

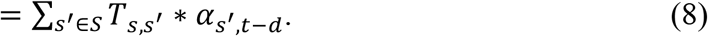

The denominator in Eq. 2, *P*(*o*_*t+1*_|*o*_*1*_, *…, o*_*t*_), is computed by conditioning on state and *ϕ*_*t*_:

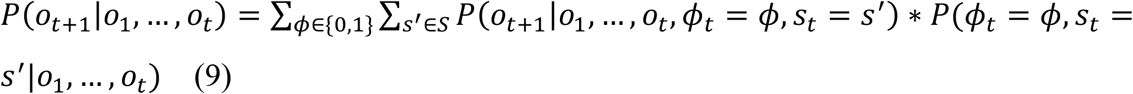

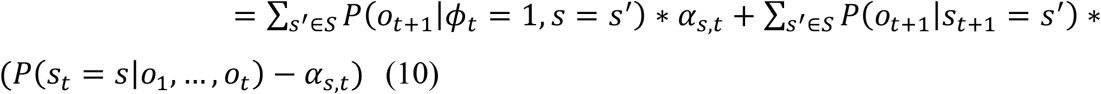

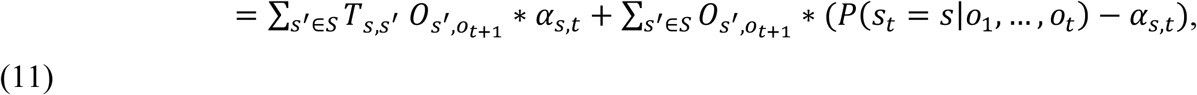

where the belief *P*(*s*_*t*_ = *s*|*o*_*1*_, *…, o*_*t*_) is computed recursively as Eq. 6 by replacing *D*_*s,d*_ with *P*(*d*_*t*_ *> d*|*s*_*t*_= *s*).

Vectorized prediction errors are gated by the probability of having just exited a state according to

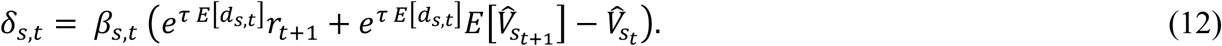

Here, the future reward, *r*_*t+1*_, and expected value of the successor state at time *t + 1*, 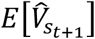 are each exponentially discounted by the expected dwell time spent in state *s* before transition, *E*[*d*_*s,t*_*A*]. To regulate the strength of temporal discounting, we used a discount factor of τ = 0.05. The expecation 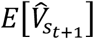 is computed by conditioning on the hypothesis that the process transitioned out of state *s* at time *t* and integrated over successor states *s*^*’*^:

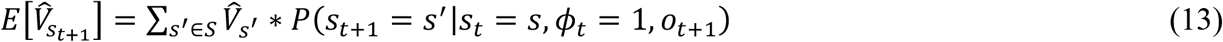

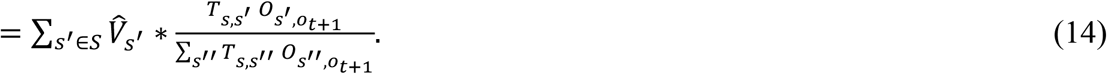

The dwell time is unknown owing to the nature of partial observability, but it can be estimated by integrating over duration spent in state *s* as for the computation of *α*. The sum is taken out until the last non-empty observation is observed since the dwell time could not be longer.

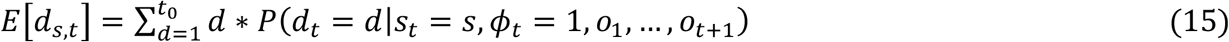

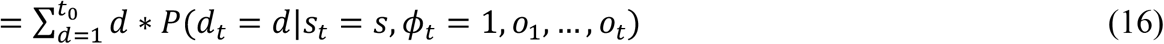

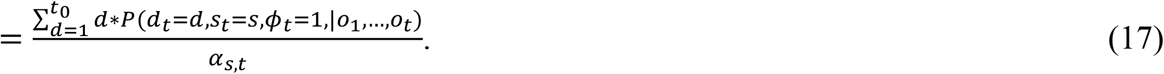

Values for each state are updated with the total prediction error over all states, ∑_*s∈S*_ *δ*_*s,t*_, at each time point,

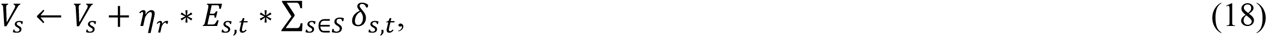

where *η*_*r*_ = 0.5 is the learning rate and controls the speed of learning the state value. The eligibility trace, *E*_*s,t*_, records the visiting of each state from the start of a trial,

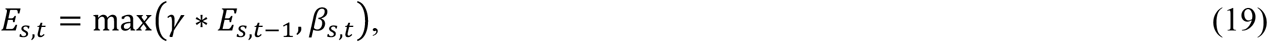

where γ = 0.95 is a temporal decay parameter, determining how far the prediction error could backpropagate to the states preceeding the current state.

When a non-empty observation is observed, the dwell time distribution is updated by a Gaussian density function that centers on the time that passed since the last non-empty observation.

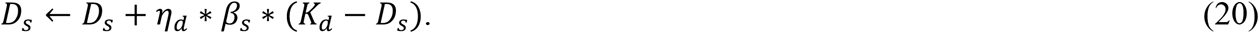

Here, *η*_*d*_ = 0.3 is the learning rate for these dwell distributions. *K*_*d*_ is a Gaussian kernel with mean *d*, and standard deviation, *CV*_*d*_ ∗ *d*, with coefficient of variation *CV*_*d*_ = 0.05. To keep *P*(*d*|*s*) non vanishing for all reasonable dwell times, we fixed the baseline probability to *D*_*b*_ = *1*0^*−4*^ for all *d*.

### Reward prediction error and neural firing

It has been suggested that dopamine neurons encode reward prediction error. We converted the total prediction error over all states, ∑_*s∈S*_ *δ*_*s,t*,_ into equivalent firing rate and compared it with the averaged firing rate of dopamine neurons:

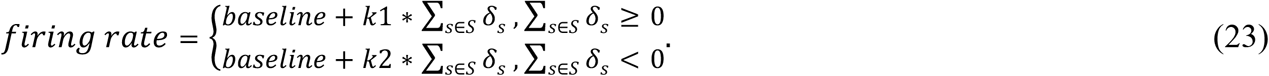

Baseline firing was set to 3Hz. *k1* = 5 and *k*2 = *−*2 were the scale factor for positive and negative prediction errors, respectively. Different scale factors were used to reflect the fact that negative errors were underrepresented in vivo.

We did not formally fit the parameters to dopamine neurons’ activities. Instead, we chose parameters manually to ensure that the reward prediction error signals quantitively match the neural activities. The results were robust and quantitively invariant across a wide range of parameter values. In addition, the results were not sensitive to the initial state value or dwell time distribution. The simulations mimicked the task event timing used in the animal experiment. To reduce any potential variance caused by the initialization and inaccurate estimation of state values and dwell distributions, we excluded the first 10 sessions from the analysis shown in the results, and all results were averaged across 20 independent simulations.

For simplicity, we did not include free-choice trials but treated them similarly to forced-trials on the same side. We assumed that the model always selected the fluid well that led to the reward, since rats exhibited high accuracy (>80%) during the forced-choice trials. Fully modeling the free-choice trials and the corresponding choice did not change any of the reported results.

### Model 1

The task state space in model 1 was designed to reflect rats’ physical location and observation when performing the task, composing seven different states, i.e., trial start, left cue, right cue, left well, right well, left reward 1, left reward 2, right reward 1, right reward 2 and inter-trial interval (Fig. 1a). The transition matrix, shown in the left panel of Fig. 4b, controlled the transitions between states. We included observations that rats indeed observed during the task, i.e., light onset, odor cues signaling the left and right choice, rewards, light offset, and a null (i.e., empty) observation (Fig. 4b, right panel). Each state brought about a non-empty observation that indicated a transition into that state. We assigned a high probability (0.95) to the only non-null observation and 0.05 to the null observation for each state. The remaining possible observations each was assigned a background probability of 10^*-4*^ to avoid the vanishing of probabilities, and the observation probabilities for each state were normalized by dividing their sum to ensure that the summation of observation probabilities for each state was 1.

We hypothesized that the HP-lesioned animals had difficulty in maintaining the hidden information after making choices, as what was observed in the OFC-lesioned animals. Therefore, we simulated an HC lesion by reducing the lesioned model’s ability to retain information when choices were made. Specifically, we set the probabilities of transitions from cue states to the well state on the same side to 0.55, and the probabilities of the transitions to the other well to 0.45 (Fig. 4b, left panel).

### Model 2

We constructed a hierarchical task space for model 2, with the lower-level describing the process happening in each trial and upper-level controlling the transition between blocks. The states in the lower-level were defined similarly to the states in model 1, except that separate states were introduced to describe the reward contingency in each block (Fig, 5a). The probabilities of transitions from the inter-trial state to each trial start states were updated with a learning rule,

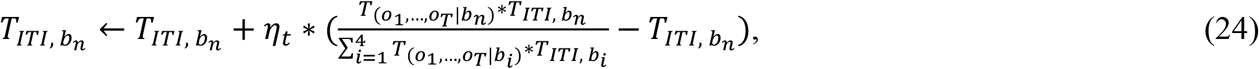

where 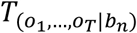 was the probability of observing the sequence of observations, *o*_*1*_, *…, o*_*T*_, in block n. The probability of transition from inter-trial interval to the block *n* trial start state, 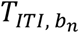, would become larger if *o*_*1*_, *…, o*_*T*_ were more likely to be observed in block n, and *η*_*t*_ controlled the learning speed. Other transition probabilities were also not updated (i.e. those describing transitions between lower-level states within a trial) since they were not changed through training.

Task observations and their probabilities were assigned similarly to those in the model 1. Unlike state dwell distributions in model 1, which were updated to reflect the changing timing of the reward, the dwell distribution in the model 2 was fixed during the training process. This was because different temporal reward contingencies were represented by separate states in model 2. As a result, estimates of state durations did not change, so we fixed dwell distributions for model 2 according to the mean delay to reward in the actual task.

Hippocampus is widely recognized as an important brain region for representing cognitive states, which reflect an agent’s location within a cognitive map. Therefore, we hypothesized that HCx rats failed to update and track their current block in a session. To simulate the effect of HC lesion, we introduced a higher variance between transitions in the top layer of the state space in our model (Fig, 5b). Specifically, 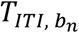 was set to 0.15 whenever it was smaller than 0.15.

## Author Contributions

YKT, TK, and GS conceived of and designed the experiments, YKT and MMC conducted the behavioral training and single unit recording, and ZZ conducted the modeling after the experiment, with advice and assistance from AJL. All authors contributed to intepreting the data and writing the manuscript.

## Acknowledgments

This work was supported by the Intramural Research Programs at the National Institute on Drug Abuse and the National Institute on Mental Health. The opinions expressed in this article are the authors’ own and do not reflect the view of the NIH/DHHS. The authors have no conflicts of interest to report.

## References

Akam, T., Costa, R., and Dayan, P. (2015). Simple plans or sophisticated habits? State, transition and learning interactions in the two-step task. PLoS Computational Biology 11(I12): e1004648. doi:10.1371/journal.pcbi.1004648.

Behrens, T.E., Muller, T.H., Whittington, J.C.R., Mark, S., Baram, A.B., Stachenfeld, K.L., and Kurth-Nelson, Z. (2018). What is a cognitive map? Organizing knowledge for flexible behavior. Neuron 100, 490–509.

Botvinick, M.M. (2012). Heirarchical reinforcement learning and decision making. Current Opinion in Neurobiology 22, 956–962.

Bromberg-Martin, E.S., Matsumoto, M., Hong, S., and Hikosaka, O. (2010). A pallidus-habenula-dopamine pathway signals inferred stimulus values. Journal of Neurophysiology 104, 1068–1076.

Costa, K.M., Scholz, R., Lloyd, K., Moreno-Castilla, P., Gardner, M.P.H., Dayan, P., and Schoenbaum, G. (2023). The role of the lateral orbitofrontal cortex in creating cognitive maps. Nature Neuroscience 26, 107–115.

Daw, N., Courville, A.C., and Touretzky, D.S. (2006). Representation and timing in theories of the dopamine system. Neural Computation 18, 1637–1677.

Dezfouli, A., and Balleine, B.W. (2012). Habits, action sequences and reinforcement learning. European Journal of Neuroscience 35, 1036–1051.

Duvelle, E., Grieves, R.M., and van der Meer, M.A. (2022). Temporal context and latent state inference in the hippocampal splitter signal. eLIFE 12, e82357.

Eichenbaum, H. (2000). Hippocampus: Mapping or memory? Current Biology 10, R785–R787.

Farovik, A., Place, R.J., McKenzie, S., Porter, B., Munro, C.E., and Eichenbaum, H. (2015). Orbitofrontal cortex encodes memories within value-based schemas and represents contexts that guide memory retrieval. Journal of Neuroscience 35, 8333–8344.

Fiorillo, C.D., Newsome, W.T., and Schultz, W. (2008). The temporal precision of reward prediction in dopamine neurons. Nature Neuroscience 11, 966–973.

Garr, E., Cheng, Y., Jeong, H., Brooke, S., Castell, L., Bal, A., Magnard, R., Namboodiri, V.M.K., and Janak, P.H. (2023). Mesostriatal dopamine is sensitive to specific cue-reward contingencies. BioRxiv. https://doi.org/10.1101/2023.06.05.543690.

Hennig, J.A., Pinto, S.A.R., Yamaguchi, T., Linderman, S.W., Uchida, N., and Gershman, S.J. (2023). Emergence of belief-like representations through reinforcement learning. BioRxiv. https://doi.org/10.1101/2023.04.04.535512.

Hollerman, J.R., and Schultz, W. (1998). Dopamine neurons report an error in the temporal prediction of reward during learning. Nature Neuroscience 1, 304–309.

Jeong, H., Taylor, A., Floeder, J.R., Lohmann, M., Mihalas, S., Wu, B., Zhou, M., Burke, D.A., and Namboodiri, V.M.K. (2022). Mesolimbic dopamine release conveys causal associations. Science 378, AOP.

Jo, Y.S., Lee, J., and Mizumori, S.J. (2013). Effects of prefrontal cortical inactivation on neural activity in the ventral tegmental area. Journal of Neuroscience 33, 8159–8171.

Kobayashi, K., and Schultz, W. (2008). Influence of reward delays on responses of dopamine neurons. Journal of Neuroscience 28, 7837–7846.

Lak, A., Nomoto, K., Keramati, M., Sakagami, M., and Kepecs, A. (2017). Midbrain dopamine neurons signal belief in choice accuracy during a perceptual decision. Current Biology 27, 821–832.

Lee, R.S., Engelhard, B., Witten, I.B., and Daw, N.D. (2022). A vector reward prediction error model explains dopaminergic heterogeneity. BioRxiv.

Matsumoto, M., and Hikosaka, O. (2009). Two types of dopamine neuron distinctly convey positive and negative motivational signals. Nature 459, 837–841.

Millidge, B., Song, Y., Lak, A., Walton, M.E., and Bogacz, R. (2023). Reward-bases: dopaminergic mechanisms for adaptive acquisition of multiple reward types. BioRxiv. https://doi.org/10.1101/2023.05.09.540067.

Mirenowicz, J., and Schultz, W. (1994). Importance of unpredictability for reward responses in primate dopamine neurons. Journal of Neurophysiology 72, 1024–1027.

Morris, G., Nevet, A., Arkadir, D., Vaadia, E., and Bergman, H. (2006). Midbrain dopamine neurons encode decisions for future action. Nature Neuroscience 9, 1057–1063.

Nomoto, K., Schultz, W. T W., and Sakagami, M. (2010). Temporally extended dopamine responses to perceptually demanding reward-predictive stimuli. Journal of Neuroscience 30, 10692–10702.

O’Keefe, J., and Nadel, L. (1978). The Hippocampus as a Cognitive Map (Clarendon Press).

Papageorgiou, G.K., Baudonnat, M., Cucca, F., and Walton, M.E. (2016). Mesolimbic dopamine encodes prediction errors in a state-dependent manner. Cell Reports 15, 221–228.

Roesch, M.R., Calu, D.J., and Schoenbaum, G. (2007). Dopamine neurons encode the better option in rats deciding between differently delayed or sized rewards. Nature Neuroscience 10, 1615–1624.

Sadacca, B.F., Jones, J.L., and Schoenbaum, G. (2016). Midbrain dopamine neurons compute inferred and cached value prediction errors in a common framework. eLIFE 5, e13665.

Schapiro, A.C., Turk-Brown, N.B., Norman, K.A., and Botvinick, M.M. (2016). Statistical learning of temporal community structure in the hippocampus. Hippocampus 26, 3–8.

Schuck, N.W., Cai, M.B., Wilson, R.C., and Niv, Y. (2016). Human orbitofrontal cortex represents a cognitive map of state space. Neuron 91, 1402–1412.

Schultz, W. (2016). Dopamine reward prediction-error signalling: a two-component response. Nature Reviews Neuroscience 17, 183–195.

Smith, D.M., and Bulkin, D.A. (2014). The form and function of hippocampal context representations. Neuroscience and Biobehavioral Reviews 40, 52–61.

Stachenfeld, K.L., Botvinick, M.M., and Gershman, S.J. (2017). The hippocampus as a predictive map. Nature Neuroscience 20, 1643–1653.

Starkweather, C.K., Babayan, B.M., Uchida, N., and Gershman, S.J. (2017). Dopamine reward prediction errors reflect hidden-state inference across time. Nature Neuroscience 20, 581–589. Starkweather, C.K., Gershman, S.J., and Uchida, N. (2018). The medial prefrontal cortex shapes dopamine reward prediction errors under state uncertainty. Neuron 98, 616–629.

Starkweather, C.K., and Uchida, N. (2021). Dopamine signals as temporal difference errors: recent advances. Current Opinion in Neurobiology 67, 95–105.

Takahashi, Y.K., Batchelor, H.M., Liu, B., Khanna, A., Morales, M., and Schoenbaum, G. (2017). Dopamine neurons respond to errors in the prediction of sensory features of expected rewards. Neuron 95, 1395–1405.

Takahashi, Y.K., Langdon, A.J., Niv, Y., and Schoenbaum, G. (2016). Temporal specificity of reward prediction errors signaled by putative dopamine neurons in rat VTA depends on ventral striatum. Neuron 91, 182–193.

Takahashi, Y.K., Roesch, M.R., Wilson, R.C., Toreson, K., O’Donnell, P., Niv, Y., and Schoenbaum, G. (2011). Expectancy-related changes in firing of dopamine neurons depend on orbitofrontal cortex. Nature Neuroscience 14, 1590–1597.

Takahashi, Y.K., Stalnaker, T.A., Mueller, L.E., Harootonian, S.K., Langdon, A.J., and Schoenbaum, G. (2023). Dopaminergic prediction errors in the ventral tegmental area reflect a multithreaded predictive model. Nature Neuroscience 26, 830–839.

Tolman, E.C. (1948). Cognitive maps in rats and men. Psychological Review 55, 189–208. Waelti, P., Dickinson, A., and Schultz, W. (2001). Dopamine responses comply with basic assumptions of formal learning theory. Nature 412, 43–48.

Wassum, K.M., Ostlund, S.B., Balleine, B.W., and Maidment, N.T. (2011). Differential dependence of Pavlovian incentive motivation and instrumental incentive learning processes on dopamine signaling. Learning & Memory 18, 475–483.

Watabe-Uchida, M., Zhu, L., Ogawa, S.K., Vamanrao, A., and Uchida, N. (2012). Whole-brain mapping of direct inputs to midbrain dopamine neurons. Neuron 74, 858–873.

Wilson, R.C., Takahashi, Y.K., Schoenbaum, G., and Niv, Y. (2014). Orbitofrontal cortex as a cognitive map of task space. Neuron 81, 267–279.

