## Supplemental Figures for "Expectancy-related changes in firing of dopamine neurons depend on hippocampus"

### Figure S1

| AP (mm) | ML (mm) | DV (mm) | Volume (nl) |
|---|---|---|---|
| -2.2 | 1.0 | -3.0 | 100 |
| -3.0 | 1.8 | -2.8 | 100 |
| -4.0 | 2.8 | -2.6 | 100 |
| -4.8 | 4.8 | -6.5 | 125 |
| -4.8 | 4.5 | -3.5 | 100 |
| -5.7 | 4.9 | -4.0 | 100 |
| -5.7 | 5.1 | -6.0 | 125 |

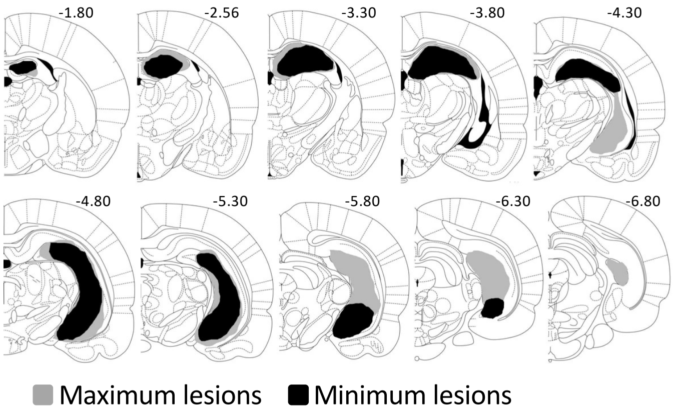


**Figure S1: Surgical coordinates and extent of ipsilateral hippocampal lesions.** Table gives volumes and coordinates (AP and ML relative to bregma and DV relative to brain surface) of injections. Brain sections illustrate the extent of the maximum (gray) and minimum (black) lesion at each level in HCx in the lesioned rats.

### Figure S2

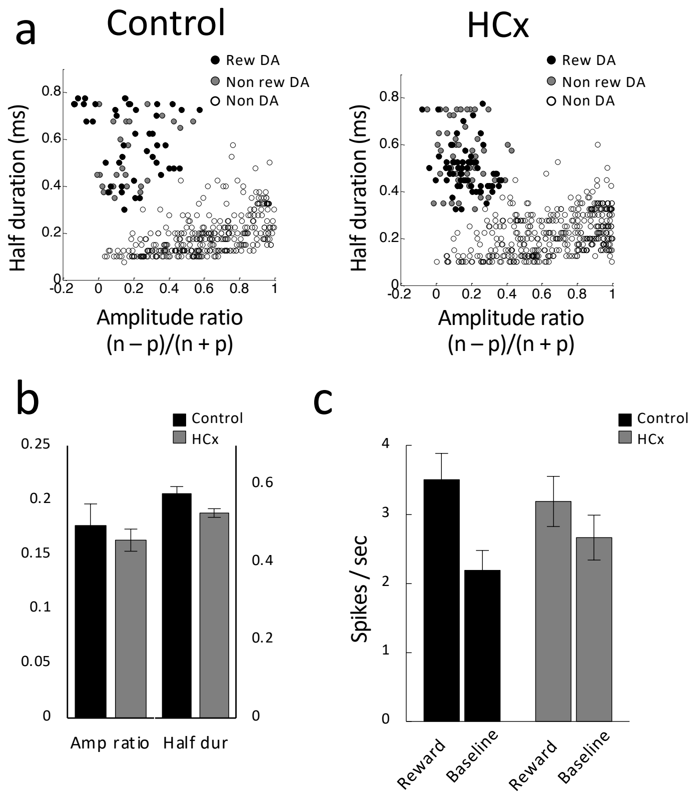


**Figure S2: Changes in reward-evoked activity of reward-responsive dopamine neurons to reward predictive odor cues. (a)** Results of cluster analysis based on the half time of the spike duration and the ratio comparing the amplitude of the first positive and negative waveform segments ((n-p)/(n+p)) in control (left) and HCx (right) groups. Reward responsive dopamine neurons (Rew DA), reward non-responsive dopamine neurons (Non rew DA), non dopamine neurons (non DA). **(b)** Bar graphs indicating average amplitude ratio and half duration of putative dopamine neurons in control (black) and HCx (gray) groups. **(c)** Average firing of putative dopamine neurons to reward and baseline in control (black) and HCx (gray) groups. Error bars, S.E.M. 2-way ANOVA comparing group (control/HCx) x epoch (reward/baseline) revealed a significant main effect on epoch ($F_{1, 108} = 91.8$, $p < 0.01$) and a significant interaction between group x epoch ($F_{1, 108} = 17.0$, $p < 0.01$).

### Figure S3

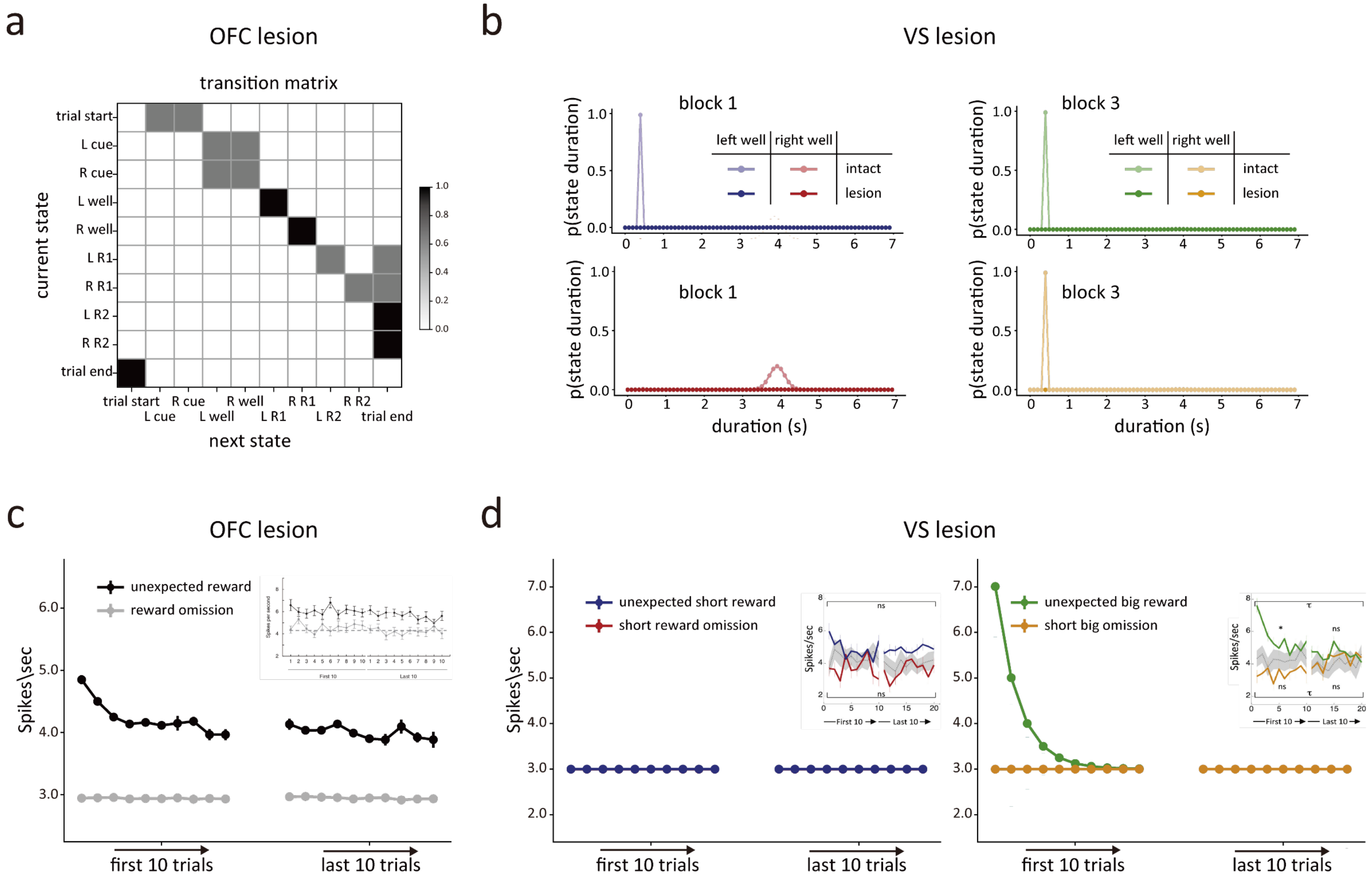


**Figure S3: Reproduce the changes in dopamine neuron firing after OFC or ventral striatal lesions. (a and c)** As indicated by (a), we fully blur the ability of the model to maintain the transition probabilities between the odor cues and the corresponding well states. By eliminating the ability to differentiate between states after actions, the model can account for the dopamine neurons response at the time of unexpected reward delivery (black lines, (c)) or omission (grey lines, (c)). Insert: Average firing of dopamine neurons after reward delivery (black lines) or omission (gray lines) in rats with OFC lesions. **(b and d)** Left panel in (b): Dwell-time distributions learned at the end of block 1 for the state left well in the short delay condition (blue) and state right well in the long delay condition (red) for the intact, and VS lesion models. Right panel in (b): Dwell-time distributions in the same format as the upper left panel, but at the end of block 3 for state left well in the big reward condition (green) and state right well in the small reward condition (orange). By preventing the model from learning and using precise duration staying in each state, the model fails to learn the dwell distribution (b), resulting in no prediction error when the short reward is delivered (blue lines, (d)) or omitted (red lines, (d)) unexpectedly, but the prediction error to the unexpected big reward delivery (green lines, (d)) and omission (yellow lines, (d)) remains intact. These results are consistent with the response patterns observed in dopamine neurons after ventral striatal lesions (inserts).
